# Pyrimidine sufficiency is required for Sae two-component system signaling in *Staphylococcus aureus*

**DOI:** 10.1101/2025.03.20.644390

**Authors:** Dennis A. DiMaggio, Won-Sik Yeo, Shaun R. Brinsmade

## Abstract

Nucleotide metabolism in pathogens is essential for their virulence, supporting growth, survival, and immune evasion during infection. Virulence in *Staphylococcus aureus* is driven by the production of virulence factors that facilitate nutrient acquisition and promote immune evasion and subversion. One key virulence regulatory system is the Sae two-component system (TCS), which upregulates the production of various virulence factors. The sensor histidine kinase SaeS, a member of the intramembrane family of histidine kinases (IM-HKs), lacks a signal-binding domain, leaving the mechanisms by which these HKs sense signals and regulate gene expression unclear. We report that de novo pyrimidine biosynthesis is essential for maintaining Sae activity. Disruption of genes involved in pyrimidine biosynthesis reduces Sae-dependent promoter activity under pyrimidine-limited conditions. Phos-tag electrophoresis confirmed that pyrimidine limitation impacts SaeS kinase activity. The effect of pyrimidine limitation on SaeS was abrogated in a strain producing only the catalytic domain, suggesting that pyrimidines regulate SaeS activity at the membrane. Additionally, defective pyrimidine biosynthesis caused membrane defects and increased incorporation of free fatty acids into the membrane. Further, providing an extracellular sink for free fatty acids restored Sae activity in these mutants. Our study highlights the interplay between nucleotide metabolism and membrane integrity in regulating virulence factor expression through signal transduction systems in pathogens.

**Importance:** Virulence is often correlated with nutrient depletion, but our understanding of this coordination is incomplete. In *S. aureus*, the Sae two-component system (Sae TCS) is a major regulator of virulence factor production and secretion, but as the sensor histidine kinase SaeS lacks an obvious domain to perceive its inducing signal, basic questions surrounding how the kinase is triggered persist. This study aims to investigate the mechanism by which pyrimidines act to promote the activity of the SaeS kinase in *S. aureus* and further expands on the importance of the roles of pyrimidines in regulating envelope biogenesis. Understanding this intersection between nucleotide metabolism and virulence regulation opens up the possibility for the development of targeted anti-virulence strategies against *S. aureus* infections.

## Introduction

*Staphylococcus aureus* is a Gram-positive, opportunistic human pathogen that transiently colonizes the skin and anterior nares of approximately 30% of the human population, with 20% being persistently colonized (1–3). Further, those that are stably colonized are at a higher risk for developing an infection (4). *S. aureus* clinical manifestations can initially present as mild to severe skin and soft tissue infections (5, 6). If left untreated, these infections can lead to more devastating invasive infections such as infective endocarditis, osteomyelitis, and sepsis (7, 8). *S. aureus* has emerged as one of the most devastating pathogens in healthcare. Although nosocomial infections of *S. aureus* are a constant threat in healthcare settings, just as concerning are infections caused by isolates circulating through the broader community (9). The relatively recent increase in community acquired infections has been attributed in part to metabolic adaptation and antimicrobial resistance. This has led to an uptick in the number of *S. aureus* related deaths over the past decade, leading to a tremendous financial burden on the US healthcare system, costing up to nearly $3 billion in annual hospitalization costs (9, 10).

The survival and success of *S. aureus* as a pathogen fundamentally depends on its ability to align its physiology to the host environment. It does so by coordinating central metabolic processes with the production of virulence factors that serve to enhance colonization while facilitating evasion from the host immune response. The combined action of central metabolism and virulence factor production contribute to the fitness of the bacterium, which is described by the capacity of the bacterium to adapt, replicate, and persist within the host environment. One of the central metabolic pathways that has emerged as a critical determinant of both bacterial fitness and virulence regulation is the *de novo* pyrimidine biosynthetic pathway (9, 11), which provides essential precursors for DNA and RNA synthesis, cell wall components, and various cellular cofactors (12). *De novo* pyrimidine biosynthesis is a highly conserved metabolic process from microbes to animals alike. Like in other organisms, *S. aureus* pyrimidine synthesis begins with the synthesis of *N-*carbamoyl-*L-*aspartate. This first committed step is catalyzed by aspartate transcarbamoylase (PyrB) and utilizes carbamoyl-phosphate, a common metabolite in pyrimidine and nitrogen metabolism. Successive transformations by dihydroorotase (PyrC), dihydroorotate dehydrogenase (PyrD), orotate phosphoribosyltransferase (PyrE), and uridine monophosphate synthase (PyrF) result in the formation of UMP (**Fig. 1A**) (13, 14). UMP can be further processed and converted into the different pyrimidine nucleosides (uridine, cytidine, and thymidine) and nucleobases (uracil, cytosine, and thymine) required for basic cellular processes (**Fig. 1A**) (13–16). The process of synthesizing pyrimidines *de novo* is regulated by the pyrimidine biosynthesis regulator, PyrR (17). PyrR is predicted to be a bifunctional protein that acts predominantly as a uridine-dependent transcriptional regulator on expression of the pyrimidine biosynthetic gene cluster, but also acts as a <u>u</u>racil <u>p</u>hospho<u>r</u>ibosyltransfer<u>ase</u> (UPRTase) (17–20). PyrR regulation occurs in a PRPP– and UMP-dependent fashion. When intracellular levels of UMP are sufficient, PyrR binds to UMP, which enhances PyrR-mRNA binding and promotes transcriptional attenuation of the pyrimidine biosynthetic gene cluster (9, 19). Further, when UMP is low and PyrR binds PRPP, there is a drop in the binding affinity of PyrR for its mRNA targets and PyrR acts as a UPRTase to increase intracellular UMP levels. This pathway must be properly regulated in *S. aureus* as pyrimidine metabolism is correlated with significant impacts the overall fitness and virulence potential of the bacterium (9, 11, 21). Although others have shown this correlation both *in vitro* and *in vivo*, the mechanism behind pyrimidine-mediated regulation of virulence remains unclear.

**Figure 1:**
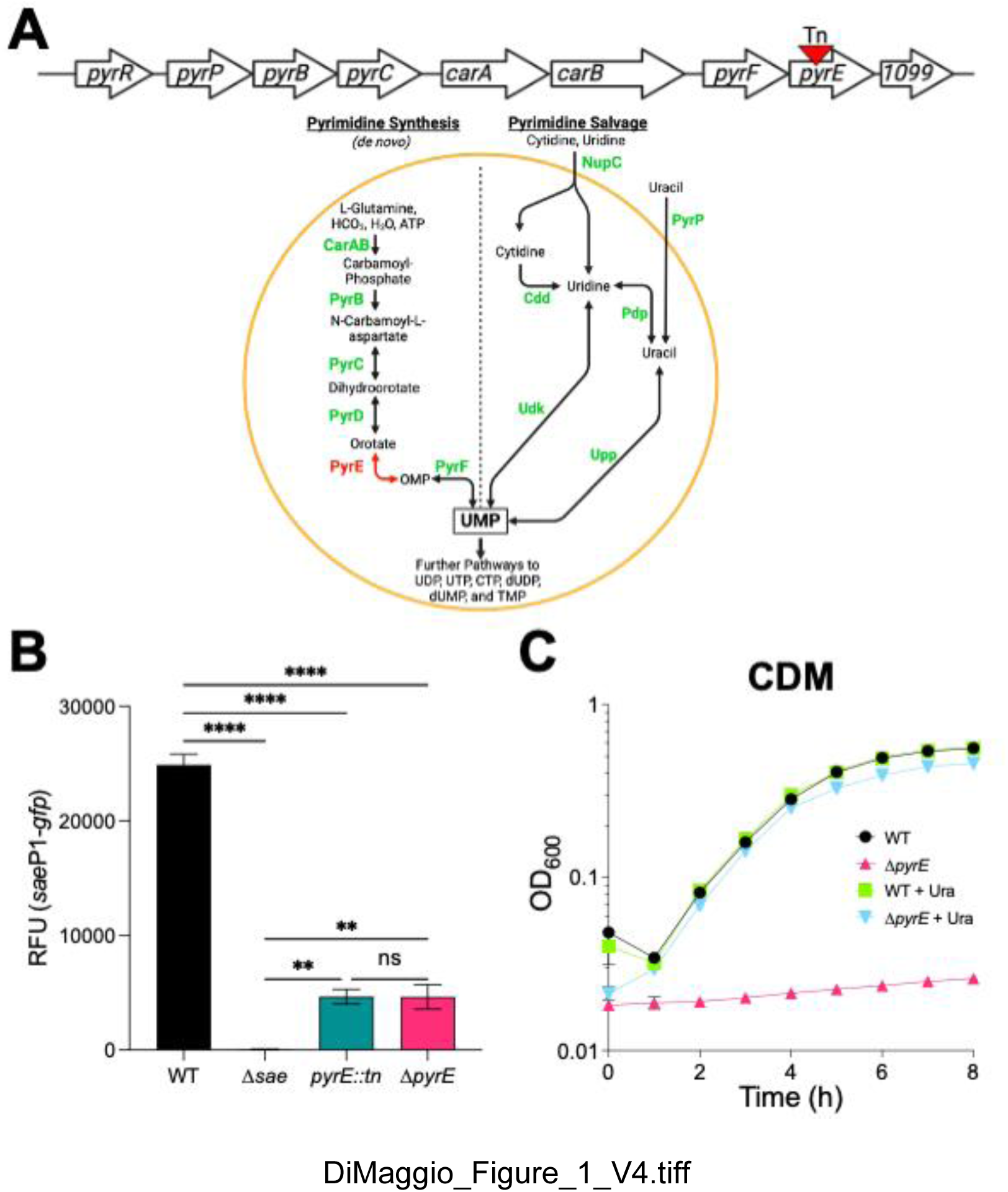
Loss of *pyrE* results in lower basal *sae*P1 promoter activity and pyrimidine auxotrophy. **(A)** (Top) Pyrimidine biosynthetic gene cluster of *S. aureus* annotated by homology to that of *Bacillus subtilis.* The transposon insertion identified in *pyrE* is indicated with an inverted red triangle. (Bottom) Current understanding of *de novo* and salvage pathways for pyrimidine biosynthesis in *S. aureus,* by analogy to the model organism *B. subtilis.* PyrR bifunctional pyrimidine regulatory protein, uracil phosphoribosyltransferase; CarAB, carbamoyl phosphate synthetase small and large subunits; PyrB, aspartate carbamoyltransferase; PyrC, dihydroorotase; PyrD, dihydroorotate dehydrogenase; PyrE orotate phosphoribosyltransferase; PyrF, orotidine 5’-phosphate decarboxylase; NupC, nucleotide permease; PyrP, uracil permease; Cdd, cytidine deaminase; Pdp, pyrimidine nucleoside phosphorylase; Udk, uridine kinase; Upp, uracil phosphoribosyltransferase (12, 14, 98) **(B)** The indicated LAC strains were grown in TSB to stationary phase (16 h), at which time *sae*P1 promoter activity was measured. Data are plotted as relative fluorescent units (RFU [GFP/OD_600_]). **(C)** Wild type (WT) and *ΔpyrE* mutant strains were grown in chemically defined medium (CDM) ± uracil supplementation (Ura; 200 µM). Growth was monitored as increase in absorbance at 600 nm (OD_600_) and plotted over time. For panels B and C, data are representative of at least three independent experiments and plotted as mean ± SEM. For panel C, error bars are plotted for all time points; in many cases they are too small to be seen. ****p<0.0001 relative to WT, One-way ANOVA with Tukey’s post-test. ns, not significant.

Production of virulence factors may incur a fitness cost. This may explain in part the fact that the expression and production of virulence factors is tightly controlled by a network of regulatory systems, including transcriptional regulators, two-component systems, and small regulatory RNAs (22–24). These systems enable *S. aureus* to occupy different niches by sensing and responding to the various environmental conditions in a coordinated manner and producing precisely the factors required for survival. Understanding the mechanisms behind virulence factor regulation may provide novel therapeutic targets for treating staphylococcal infections. A growing body of research suggest so-called anti-virulence strategies offer an opportunity to mitigate the damage *S. aureus* can cause by lowering or blocking the ability of the cells to produce virulence factors (25–27).

One major regulator of *S. aureus* virulence is the SaeRS two-component system (TCS). This TCS is responsible for regulating over 20 different secreted and surface-associated virulence factors that facilitate nutrient acquisition and assist in evading and subverting the immune response during infection (28). The *sae* (*<u>S</u>. <u>a</u>ureus* <u>e</u>xoprotein expression) locus was discovered in 1994 by Giraudo *et al.* while characterizing a transposon mutant defective in exoprotein secretion (29–32). This locus consists of an operon containing the four *sae* genes, *saeP, saeQ, saeR, and saeS. saeR* encodes the DNA-binding response regulator (RR) protein, and *saeS* encodes the membrane-spanning sensor histidine kinase (HK) (28). *saeP* and *saeQ* encode auxiliary proteins that have been shown to stimulate phosphatase activity of SaeS toward SaeR∼P, resetting the TCS to its pre-activation state (33). The expression of these genes is driven by two promoters, P1 and P3. The P1 promoter is inducible and controls the transcription of the entire *sae* operon, while the P3 promoter located in *saeQ* drives constitutive expression of *saeR* and *saeS* (34). SaeS activity is triggered by neutrophil-produced factors, most notably human neutrophil peptides (HNP1-3) by an unknown mechanism (35). In response, SaeS autophosphorylates its conserved histidine residue. Then, SaeR is phosphorylated using the phosphohistidine of SaeS as a substrate, resulting in signal transduction (36, 37). This generates SaeR∼P, which binds to site-specific sequences to promote transcription of the target genes (38, 39). One target of SaeR∼P is the *sae*P1 promoter. This creates a positive feedback loop, resulting in signal amplification and preventing Sae activity from becoming excessive.

In canonical TCSs, the HK is capable of receiving input signals through a large extracytoplasmic input domain. In contrast, SaeS lacks this domain, and the extracellular-facing portion of the protein comprises only a 9 amino acid linker (40). As a result, SaeS is nontraditional sensor HK and is considered a member of the intramembrane family of sensor histidine kinases (IM-HK) (41). How SaeS responds to HNPs is an open question. Even at a more foundational level, our knowledge of how IM-HKs like SaeS function is incomplete (42–44). Herein, we report that disrupting *de novo* pyrimidine biosynthesis reduces Sae-dependent promoter activity under pyrimidine-limited conditions. Using PhosTag electrophoresis we determine that pyrimidine limitation reduces SaeS kinase activity. This effect depends on the N-terminal membrane-spanning domain of SaeS, suggesting that pyrimidines act at the membrane to promote Sae kinase activity. Consistent with known roles of pyrimidine nucleoside triphosphates in cell membrane and envelope biogenesis, we show that pyrimidine limitation results in a thicker cell wall, irregular membrane, and an accumulation of free fatty acids. Removal of free fatty acids by providing an extracellular sink for these lipids is sufficient to restore Sae activity during pyrimidine limitation. The ripple effects generated by defects in pyrimidine metabolism highlight the critical roles that central metabolic pathways play in maintaining cellular homeostasis and virulence potential in *S. aureus*, making pyrimidine biosynthesis a promising target for the development of novel anti-virulent therapeutics.

## Results

### Disruption of *pyrE* reduces basal Sae TCS activity and pyrimidine auxotrophy

As a member of the IM family, little is known how SaeS perceives its signals. To gain insight on how SaeS functions mechanistically, we conducted a transposon mutagenesis screen using a pMA15 plasmid system (45) to identify factors required for SaeS kinase activity. Using this screen, we isolated a transposon insertion in the gene encoding orotate phosphoribosyltransferase (*pyrE*), a critical enzyme for *de novo* pyrimidine biosynthesis (**Fig. 1A**). To indirectly measure Sae activity, we introduced a Sae-dependent transcriptional reporter (*sae*P1-*gfp*) into *S. aureus* strains and grew the cells to stationary phase (16 h) in rich, complex medium (tryptic soy broth; TSB). As expected, we detected essentially no promoter activity in the Δ*sae* mutant. We measured a 4-fold drop in promoter activity in the *pyrE::tn* mutant relative to the parent strain (**Fig. 1B**). To confirm that the *sae* defect was associated with disruption of *pyrE* and to eliminate the possibility of polar effects on SAUSA300_1099 immediately downstream (annotated to encode YqzH; function unknown; depicted as *1099* in **Fig. 1A**), we generated a Δ*pyrE* in-frame deletion mutant. The Δ*pyrE* mutant also showed the same reduction in Sae activity (**Fig. 1B**). We note that the Sae system is still capable of being activated by human neutrophil peptides (**Fig. S1**). Consistent with what is known about the role of PyrE in the *de novo* pyrimidine biosynthetic pathway in other Gram-positive and Gram-negative genera, the Δ*pyrE* mutant is a pyrimidine auxotroph and is unable to grow in chemically defined medium (CDM) unless exogenous pyrimidines like uracil are provided (**Fig. 1A and C**) (11–15, 46). Notably, the *ΔpyrE* mutant exhibited a significant growth yield defect in addition to the Sae defect when grown in TSB (**Fig. S2**). This strongly suggests that pyrimidines become limiting in this medium and indicates that *S. aureus* relies on the *de novo* pathway to support growth in TSB. As expected, pyrimidine auxotrophy (or limitation) can be chemically complemented with the addition of exogenous uracil in both CDM and TSB circumventing the need for *de novo* synthesis (**Fig. 1A, 1C, and S2**). Overall, these data suggest a link between pyrimidine sufficiency and basal Sae activity.

### Pyrimidine limitation leads to a reduction in basal SaeS kinase activity

As mentioned previously, pyrimidines are either synthesized through the *de novo* pathway as needed or taken into the cell through the pyrimidine salvage pathway to support growth. Disrupting *de novo* pyrimidine biosynthesis can result in a buildup of pyrimidine metabolite precursors and alter expression of the genes predicted to be involved in the *de novo* and salvage pathways (14, 15). While characterizing the effects of pyrimidine metabolism on regulation of the Sae TCS in *S. aureus*, we noticed that the Δ*pyrE* mutant accumulates white, crystalline material extracellularly when grown on TSB agar plates (data not shown). Further, Δ*pyrE* mutant strains of other bacterial species accumulate orotate crystals extracellularly, and in some species this is linked to defects in virulence factor production (14, 15). Therefore, we hypothesized that a pyrimidine-specific metabolite, when it accumulates intracellularly or extracellularly, reduces Sae TCS activity. To test this hypothesis, we constructed additional mutations along the *de novo* pathway and measured *sae*P1 promoter activity as a proxy for Sae activity. Interestingly, when we deleted the genes annotated to encode dihydroorotate dehydrogenase (*pyrD*) and dihydroorotase (*pyrC*), we measured essentially the same low *sae*P1*-gfp* promoter activity as that measured in the *pyrE* mutant strains (**Fig. 1B and 2A**). The Sae activity defect in these strains could be complemented chemically by supplementing exogenous uracil or cytidine, or genetically in the Δ*pyrD* mutant by ectopically expressing a wild-type copy of *pyrD^+^*under control of its native promoter (**Fig. 2A and S3)**. Although we could have tested the effects of mutations further up the *de novo* pathway, we opted not to as to avoid further interference with other biosynthetic pathways (47). Since we observed an identical reduction in Sae activity in the *ΔpyrD* and *ΔpyrC* mutant cells relative to the *ΔpyrE* mutant cells, we then reasoned that the pyrimidine-related Sae defect is due to limitation. If so, Sae activity should be unaffected during exponential phase prior to the onset of pyrimidine limitation. To test this, we collected cells that were growing exponentially in TSB (2 h) and monitored Sae activity using *sae*P1-*gfp*. Consistent with our complementation data, both the *pyrE::tn* and Δ*pyrE* mutants showed no Sae defect when levels of pyrimidine in the medium were not limiting (**Fig. 2B**). Taken together, our data suggest that the defect in basal Sae activity is not due to buildup of metabolites, but rather due to pyrimidine limitation itself.

**Figure 2:**
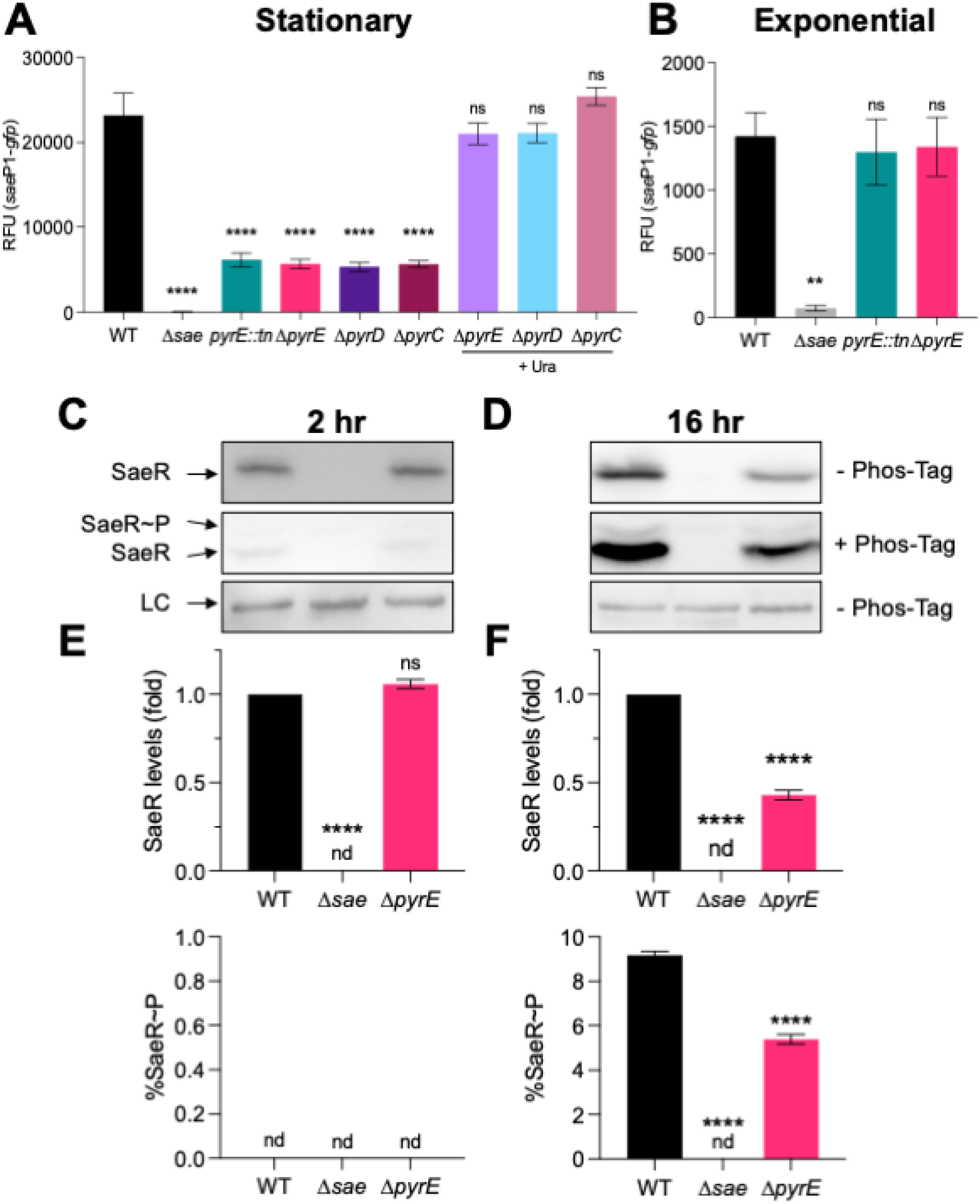
Pyrimidine limitation reduces SaeS kinase activity. The indicated strains carrying the *sae*P1-*gfp* reporter were grown in TSB ± uracil supplementation (Ura; 200 µM) to **(A)** stationary phase (16 h) and **(B)** exponential phase (2 h), at which time *sae*P1 promoter activity was measured. **(C-H)** Measurement of intracellular SaeR species. The indicated strains were grown in TSB to exponential phase (2 h) and stationary phases (16 h), at which times cell pellets were collected, normalized by OD_600_, and cell lysates then subjected to Western Blot analysis. **(C and D)** Representative Western blots (top and bottom) and Phos-tag Western blots (middle) of cell lysates using antibodies recognizing SaeR. **(E and F)** Analysis of SaeR species using Multi Gauge software (FujiFilm) showing SaeR levels relative to WT in each condition (Top), and the percentage of SaeR∼P relative to the total amount of SaeR in each sample (bottom). All data are from at least three independent experiments and are plotted as mean ± SEM. ****p<0.0001, **p<0.01,*p<0.05; One-way ANOVA with Dunnet’s post-test relative to respective WT. LC, loading control; nd, below detectible limit; ns, not significant.

Unphosphorylated SaeR cannot bind to target promoters to stimulate gene transcription (39). Only activated SaeR (SaeR∼P) can bind. Because there is little to no crosstalk between HKs and RRs, Sae output (i.e., target gene promoter activity) is solely dependent on the activity of SaeS (48). To test the hypothesis that pyrimidine limitation affects SaeS kinase activity directly, we used Phos-Tag electrophoresis (49) and Western blotting with polyclonal antibodies raised against SaeR to measure relative levels of SaeR SaeR species in Δ*pyrE* mutant cells during pyrimidine-limited and pyrimidine-replete conditions. During exponential phase, we detected slightly higher levels of SaeR in the *ΔpyrE* mutant relative to WT, but we were unable to detect SaeR∼P in either sample. This is consistent with previous studies reporting very low levels of Sae activity during this period (50, 51). Notably, we measured a general reduction in total SaeR protein levels and a concomitant drop in the fraction of SaeR∼P in the *ΔpyrE* mutant cells during pyrimidine limitation (**Fig. 2 C-F**). Given that SaeR levels are dependent in part on the activity of SaeS through SaeR∼P-mediated activation of the P1 promoter, this may confound our analysis, giving us a false sense that SaeR∼P levels are lower in the Δ*pyrE* mutant. To compare SaeS kinase activity directly without confounding autoregulation, we complemented Δ*sae* and *Δsae ΔpyrE* mutant strains with either *saeRS* (+*RS*) ectopically expressed under the control of the constitutive P3 promoter or the empty vector (+EV) and remeasured SaeR∼P as a percentage of the total SaeR in these cells. As expected, we measured no difference in total SaeR levels in Δ*pyrE* mutant cell lysates compared to WT cell lysates. Notably, we observed a dramatic reduction in the fraction of SaeR∼P in the Δ*pyrE* mutant cell lysates (**Fig. 3A-C**). Taken together, these data indicate that pyrimidine limitation results in a reduction of SaeS kinase activity.

**Figure 3:**
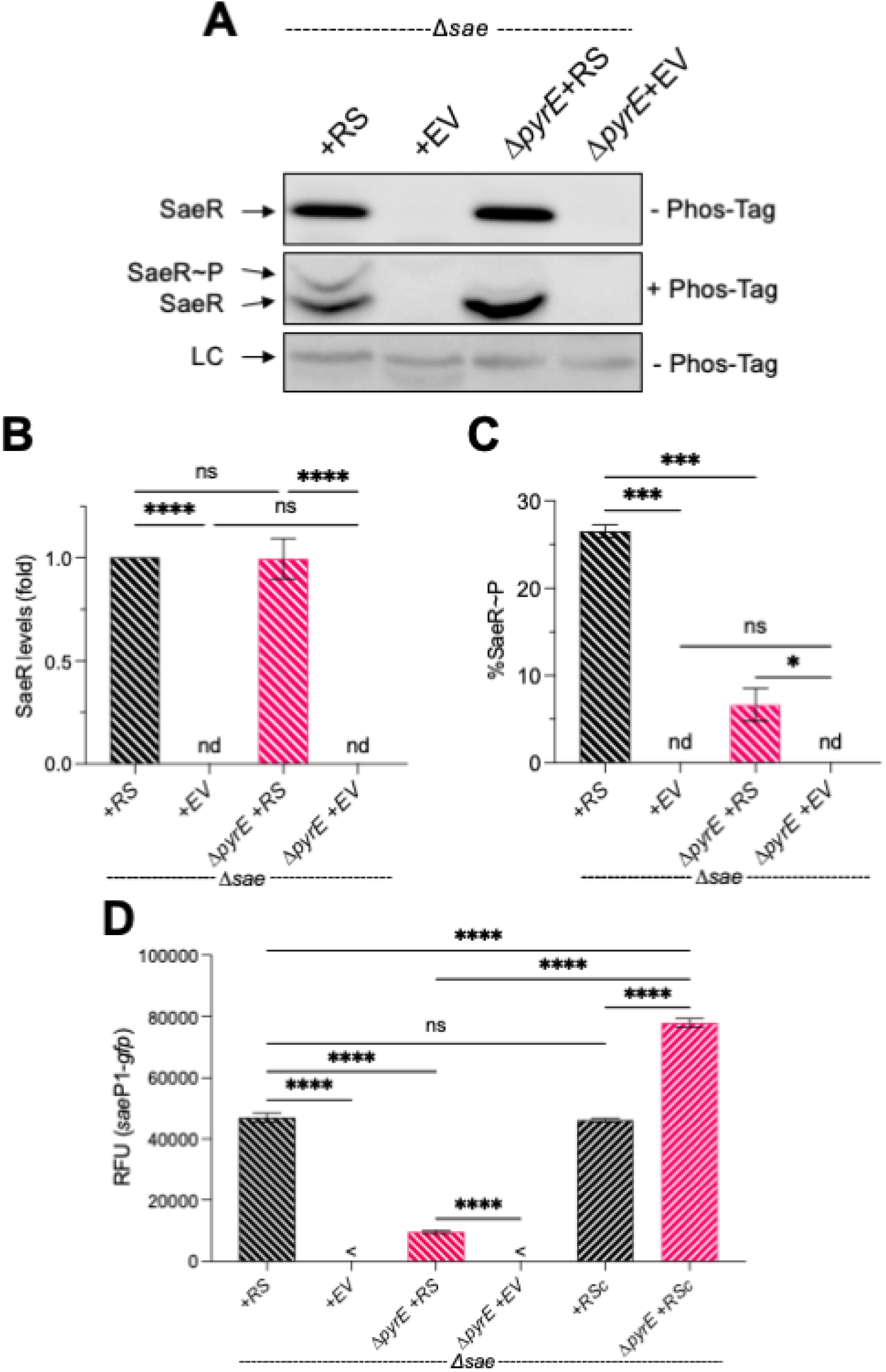
The effect of pyrimidine limitation is independent of autoregulation and acts at the membrane. (**A-C**) SaeR species present in SaePQRS-deficient strains (Δ*sae*) with either *saeRS* complemented back under control of the constitutive P3 promoter (+RS) or the empty vector (+EV) grown in TSB to stationary phase (16 h). Cell pellets were collected, normalized by OD_600_, and lysates subsequently subjected to Western Blot analysis using polyacrylamide gels ± Phos-tag acrylamide. **(A)** Representative blots are shown. **(B and C)** Densitometry analysis was performed, quantifying levels of the different SaeR species present in each sample. LC, loading control; nd, below detectible limit. **(D)** Measurement of *sae*P1-*gfp* promoter activity in stationary phase cultures (16 h) with either empty vector (EV), full length SaeS (+*RS*), or the cytosolic, catalytic domain of SaeS (+*RSc*). <, fluorescence was undetectable relative to blank containing PBS. For all panels, data are plotted as mean ± SEM, n≥3. ****p<0.0001, ***p<0.001, *p<0.05, one-way ANOVA with Tukey’s post-test. ns, not significant.

### The N-terminal, membrane-spanning domain of SaeS is the site of regulation of the Sae TCS by pyrimidines

The prevailing view is that the overall conformation of the N-terminal domain is critical for controlling kinase activity. To test if the N-terminal domain hinders the kinase activity of SaeS in Δ*pyrE* mutant cells during pyrimidine limitation, we measured SaeS kinase activity in strains producing either full-length SaeS or just the catalytic, C-terminal kinase domain located in the cytosol (SaeSc). We found that the *sae*P1*-gfp* fusion was no longer affected by pyrimidine limitation. In fact, we noted a significant increase in promoter activity compared to that measured in otherwise WT cells producing full length SaeS when SaeSc is produced (**Fig. 3D)**. Consistent with this observation, overall SaeR levels were increased in addition to the ratio of SaeR∼P/SaeR species in the Δ*pyrE* mutant during limitation (**Fig. S4**). These data indicate that pyrimidine limitation specifically targets the N-terminal membrane-spanning portion of SaeS, reducing kinase activity.

### Pyrimidine limitation significantly alters the transcriptome of *ΔpyrE* mutant cells

Given that the Sae defect in the *ΔpyrE* mutant cells during pyrimidine limitation is associated with the N-terminal membrane-spanning domain of SaeS, we wondered if pyrimidine limitation is generally linked to a defect in the envelope environment. If this is true, then the effect of pyrimidine limitation likely extends beyond Sae to many proteins in the membrane, several of which are kinases for other TCSs. To determine the full extent to which pyrimidine limitation alters gene expression and help inform a potential mechanism for Sae activation, we performed transcriptional profiling in wild-type and isogenic *ΔpyrE* mutant cells grown to early stationary phase (i.e., pyrimidine-limiting conditions). Overall, we noticed that the transcriptome was dramatically altered in the *ΔpyrE* mutant cells during limitation, with 701 genes significantly downregulated and 719 genes significantly upregulated (Log_2_ fold change ≥ 1 or ≤ –1 and *p*<0.05; *ΔpyrE*/WT) (**Table S1**). This dramatic shift in the transcriptional profile of the *ΔpyrE* mutant likely stems from the interconnectedness of pyrimidine metabolism with other central metabolic pathways and various regulators in the cell (13, 52). To determine whether pyrimidine limitation affects specific pathways preferentially, we performed KEGG enrichment analysis on the RNA-seq output using KEGGREST with the USA300_FPR3757 KEGG reference genome (KEGG;saa) (53, 54). Based on KEGG enrichment, the majority of differentially expressed genes (DEGs) annotated to be involved in 80 different KEGG pathways were downregulated in the *ΔpyrE* mutant cells during pyrimidine limitation (**Table S2**). This includes directly regulated pathways such as purine metabolism, various pathways involved in cell envelope biogenesis (e.g., fatty acid biosynthesis, teichoic acid biosynthesis, and peptidoglycan biosynthesis), and the various sugar related metabolic pathways. Further, based on this analysis, it indicates that 46 out of the 53 DEGs encoding proteins annotated to be involved in TCS signaling are downregulated in the *ΔpyrE* mutant relative to WT during pyrimidine limitation. Since the KEGG pathways base gene annotation off of functionally similar genes in previously reported systems (55), this may lead to overestimation of the effect on TCSs and prompts us to look closer into the pyrimidine limitation-induced impact on the various TCSs. To begin to answer this question, we took advantage of the fact that TCSs exhibit feedback autoregulation of the HK and RR to indirectly assess the overall impact on kinase activity of the other 15 TCSs (56, 57). We saw that, in addition to the log_2_-fold drop in *saeRS* transcripts, there was a general reduction in the expression of the other TCS coding genes in the *ΔpyrE* mutant cells during pyrimidine limitation (**Table 1**). Interestingly, the Agr quorum sensing TCS had the highest reduction in expression relative to the other systems with a 5-to 6-fold drop in *agrCA* transcript abundance.

**Table 1:** Pyrimidine limitation negatively effects expression of most two-component systems in the *ΔpyrE* mutant. RNA-seq analysis results quantifying the fold-change in expression of TCS HK and RR genes in the *ΔpyrE* mutant relative to the WT after 8 h of growth in TSB (pyrimidine-limiting condition for *ΔpyrE* mutant cells). Raw reads were trimmed using Trimmomatic. Trimmed reads were aligned and analyzed based on the reference genome (USA300 FPR3757; NC007793) using Rockhopper. p-values were determined using a Negative Binomial distribution and adjusted p-values were determined using the Benjamin Hochberg procedure built into the script. The full data set can be found in **Table S1**.

| Gene | Log<sub>2</sub> Fold Change<br>(ΔpyrEWT) | -log<sub>10</sub>p | Adjusted<br>p-value |
| --- | --- | --- | --- |
| *saeS* | -2.63 | 6.8E-19 | 0.00 |
| *saeR* | -2.07 | 7.8E-07 | 0.00 |
| *agrC* | -5.03 | 1.2E-307 | 0.00 |
| *agrA* | -6.15 | 1.2E-307 | 0.00 |
| *airS* | 1.35 | 9.8E-08 | 0.00 |
| *airR* | 0.24 | 5.6E-01 | 1.00 |
| *vraS* | -1.08 | 1.2E-04 | 0.00 |
| *vraR* | -1.70 | 2.0E-15 | 0.00 |
| *phoR* | -2.18 | 5.0E-16 | 0.00 |
| *phoP* | -2.06 | 2.7E-16 | 0.00 |
| *desK* | -0.36 | 3.4E-01 | 1.00 |
| *desR* | -0.85 | 1.4E-03 | 0.00 |
| *arlS* | -1.71 | 1.6E-10 | 0.00 |
| *arlR* | -0.87 | 4.5E-03 | 0.01 |
| *braS* | -1.21 | 7.8E-05 | 0.00 |
| *braR* | -0.25 | 4.8E-01 | 0.87 |
| *graS* | -1.25 | 1.5E-06 | 1.00 |
| *graR* | -0.16 | 6.0E-01 | 1.00 |
| *hptS* | -1.39 | 1.8E-10 | 1.00 |
| *hptR* | -2.32 | 2.1E-45 | 1.00 |
| *hssS* | 0.44 | 1.0E-01 | 1.00 |
| *hssR* | 2.06 | 1.4E-14 | 1.00 |
| *kdpD* | -3.17 | 1.4E-161 | 1.00 |
| *kdpE* | -3.22 | 3.9E-187 | 1.00 |
| *lytS* | 0.78 | 7.5E-03 | 1.00 |
| *lytR* | 0.00 | 9.7E-01 | 1.00 |
| *nreB* | -1.90 | 1.9E-14 | 0.00 |
| *nreC* | -2.40 | 4.5E-44 | 0.00 |
| *srrB* | 0.23 | 3.0E-01 | 0.58 |
| *srrA* | -0.56 | 4.5E-02 | 0.10 |
| *walK* | -0.65 | 3.7E-02 | 0.08 |
| *walR* | 0.84 | 4.3E-03 | 0.01 |

Since AgrC has a lower affinity for ATP (58) and ATP is required to activate the sensor kinases of the TCSs, we wondered if the reduction in activity of these TCSs was potentially correlated with a drop in intracellular ATP pools due to a concomitant reduction of purine biosynthesis in the *ΔpyrE* mutant cells during pyrimidine limitation. Although we did not test the effect of pyrimidine limitation on purine biosynthesis specifically by measuring ATP pools, KEGG analysis identified that 21 of the 29 DEGs were downregulated for purine biosynthesis. Further, 10 of the 14 genes previously annotated to be involved in the *de novo* purine biosynthetic pathway and all of the genes annotated for the different subunits of the ATP synthase were significantly downregulated in the *ΔpyrE* mutant during pyrimidine limitation (**Table 2 and S2**). This suggests that the cell is reducing purine production in response to pyrimidine limitation to maintain balance within the cell. In addition, many of the genes annotated to be involved in nucleotide salvage were upregulated in the Δ*pyrE* mutant cells (**Table 2**). Collectively, these data show that defective pyrimidine metabolism results in a global change to the transcriptional profile of *S. aureus* when pyrimidines are limited, including those involved in the synthesis of envelope components. Further, transcriptomic analysis suggests that pyrimidine limitation generally reduces the kinase activity of the majority of TCSs in *ΔpyrE* mutant cells.

**Table 2:** Pyrimidine limitation results in differential expression of genes involved in purine and pyrimidine biosynthesis in the *ΔpyrE* mutant. RNA-seq analysis results quantifying the fold-change in expression of annotated purine and pyrimidine biosynthesis genes as well as the annotated ATP synthase genes in the *ΔpyrE* mutant relative to the WT after 8 h of growth in TSB (pyrimidine-limiting condition for *ΔpyrE* mutant cells). Raw reads were trimmed using Trimmomatic, trimmed reads were aligned and analyzed based on the reference genome (USA300 FPR3757; NC007793) using Rockhopper. p-values were determined using a Negative Binomial distribution and adjusted p-values were determined using the Benjamin Hochberg procedure built into the script. The full data set can be found in **Table S1**.

| Pyrimidine Biosynthesis and Salvage |  |  |  |
| --- | --- | --- | --- |
| Gene | Log<sub>2</sub> Fold Change<br>(ΔpyrE/WT) | -log<sub>10</sub>p | Adjusted<br>p-value |
| *nupC* | 1.12 | 3.08E-04 | 0.00 |
| *pyrR* | 4.47 | 1.15E-307 | 0.00 |
| *pyrP* | 3.82 | 9.07E-306 | 0.00 |
| *pyrB* | 1.91 | 1.50E-16 | 0.00 |
| *pyrC* | 0.69 | 1.56E-01 | 0.32 |
| *carA* | 0.16 | 5.38E-01 | 0.95 |
| *carB* | -0.65 | 1.48E-01 | 0.30 |
| *pyrF* | -1.23 | 8.51E-05 | 0.00 |
| *pyrE* | ---- | 1.15E-307 | 0.00 |
| *pyrH* | 1.63 | 1.49E-10 | 0.00 |
| *cdd* | -0.80 | 5.79E-03 | 0.01 |
| *udk* | 0.99 | 7.62E-04 | 0.00 |
| *upp* | -2.00 | 1.02E-12 | 0.00 |
| *pyrG* | -0.70 | 6.45E-02 | 0.14 |
| *pdp* | -1.30 | 1.49E-04 | 0.00 |
| *pyrD* | 3.79 | 1.15E-307 | 0.00 |
| Purine Biosynthesis and ATP Synthesis |  |  |  |
| Gene | Log<sub>2</sub> Fold Change<br>(ΔpyrE/WT) | -log<sub>10</sub>p | Adjusted<br>p-value |
| *purA* | -0.31 | 5.83E-01 | 1.00 |
| *purR* | -0.61 | 4.38E-02 | 0.10 |
| *purE* | -1.16 | 8.60E-05 | 0.00 |
| *purK* | -2.56 | 4.73E-32 | 0.00 |
| *purC* | -3.63 | 2.08E-255 | 0.00 |
| *purS* | -4.27 | 1.15E-307 | 0.00 |
| *purQ* | -5.37 | 1.15E-307 | 0.00 |
| *purL* | -6.91 | 1.15E-307 | 0.00 |
| *purF* | -7.76 | 1.15E-307 | 0.00 |
| *purM* | -8.16 | 1.15E-307 | 0.00 |
| *purN* | -8.63 | 1.15E-307 | 0.00 |
| *purH* | -7.43 | 1.15E-307 | 0.00 |
| *purD* | -7.51 | 1.15E-307 | 0.00 |
| *purB* | 0.34 | 4.57E-01 | 0.83 |
| *atpC* | -5.10 | 1.15E-307 | 0.00 |
| *atpD* | -4.83 | 1.15E-307 | 0.00 |
| *atpG* | -4.99 | 1.15E-307 | 0.00 |
| *atpA* | -4.38 | 1.15E-307 | 0.00 |
| *atpH* | -3.86 | 1.15E-307 | 0.00 |
| *atpF* | -3.48 | 1.27E-185 | 0.00 |
| *atpA* | -4.38 | 1.15E-307 | 0.00 |
| *atpE* | -2.59 | 1.27E-46 | 0.00 |
| *atpB* | -1.87 | 1.04E-09 | 0.00 |

### Pyrimidine limitation is associated with cell wall thickening and an unstable membrane

Up to this point, we have shown that SaeS integrates pyrimidine sufficiency to regulate basal kinase activity via the membrane-spanning domain. However, how pyrimidine sufficiency is perceived by SaeS mechanistically remains unknown. Because pyrimidine nucleotides are used to activate precursors for peptidoglycan, teichoic acid, and phospholipid synthesis and that genes that direct the synthesis of these membrane constituents are differentially expressed, we reasoned that during pyrimidine limitation a modified cell envelope might contribute to the defect in SaeS kinase activity. We determined the minimum inhibitory concentration (MIC) of the *ΔpyrE* mutant and the WT parent strain for available molecules that target synthesis of major cell envelope components (59). We measured no change in susceptibility to fosfomycin (first committed step in peptidoglycan synthesis), tunicamycin (wall teichoic acid synthesis), or oxacillin (peptidoglycan crosslinking) (**Table S3**) (59–62). In contrast, we observed a 4-to 8-fold increase in the susceptibility to congo red, with a change in MIC from 0.25% in WT cells to 0.06%-0.03 in Δ*pyrE* mutant cells. Congo red was previously reported to be an inhibitor of LtaS (lipoteichoic acid [LTA] synthase) that is critical for LTA assembly (63). To better understand the change in susceptibility, we extracted lipoteichoic acids from WT and *ΔpyrE* mutant cells and found that the Δ*pyrE* mutant has increased production of LTAs (**Fig. S5AB**). Previously, uncontrolled LTA synthesis in the absence of the small membrane regulatory protein MspA was shown to result in virulence defects (64, 65). To determine whether pyrimidine limitation acts in the same regulatory pathway as MspA we performed a genetic epistasis experiment. The *mspA::tn* single mutant showed essentially the same SaeS kinase activity defect as the Δ*pyrE* mutant, as judged by phos-tag experiments. No further reduction in kinase activity was measured in the Δ*pyrE mspA::tn* double mutant. The kinase defect was complemented when a wild-type copy of *mspA^+^* was provided *in trans* in the *mspA::tn* single mutant, but not in the Δ*pyrE mspA::tn* double mutant, suggesting that the effects on Sae activity are independent from one another (**Fig. S5C**).

To get a better understanding on the impact of pyrimidine limitation on the cell envelope, we performed transmission electron microscopy (TEM) on WT, *ΔpyrE*, and *ΔpyrE* mutant cells fed exogenous uracil. Multiple observations are consistent with an abnormal cell envelope. First, Δ*pyrE* mutant cells experiencing pyrimidine limitation are nearly twice as large as WT cells; this phenotype is complemented when the missing nutrient is provided (**Fig. S6 A-D**). Second, Δ*pyrE* mutant cells under limitation conditions have a much thicker cell envelope and a visibly crenulated membrane architecture relative to WT or the chemically complemented cells (**Fig. 4A-D**). Increased envelope thickness in the *ΔpyrE* mutant is consistent with the idea that defects in central metabolism and/or increased LTA biosynthesis result in thickening of the cell envelope (11, 61, 64). In addition to the dramatic difference in membrane morphology between WT and pyrimidine-limited (*ΔpyrE* mutant) cells, *ΔpyrE* mutant cells accumulated more of the eponymous staphyloxanthin pigment compared to wild-type cells (**Fig. S7A**).

**Figure 4:**
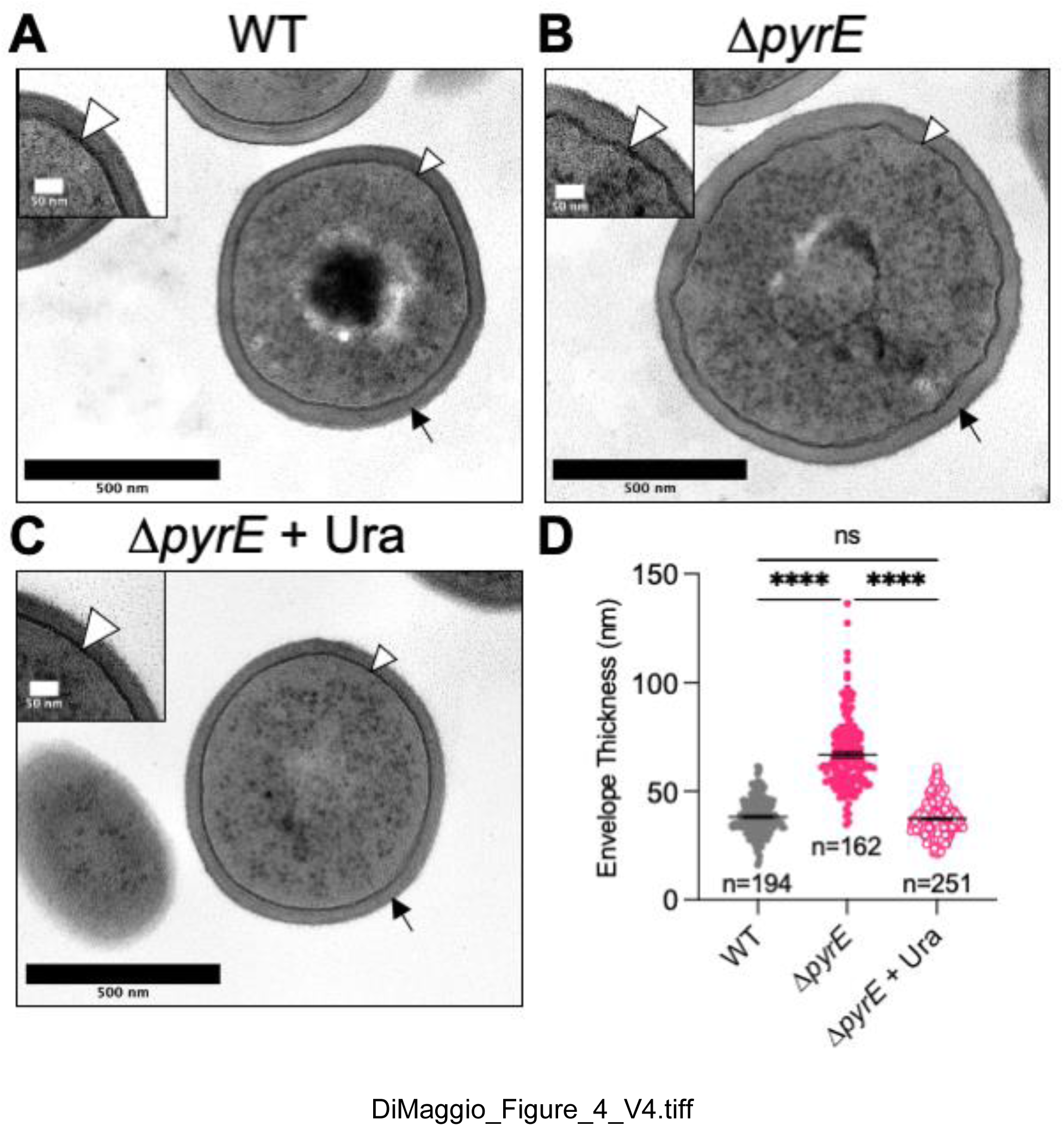
Δ*pyrE* mutant cells have a thickened cell wall and a crenulated membrane. Representative TEM images of the **(A)** wild-type (WT), **(B)** *ΔpyrE* mutant, and **(C)** *ΔpyrE* mutant chemically complemented with uracil (Ura; 200 µM). Inset images emphasize the crenulated cell membrane architecture of the Δ*pyrE* mutant. **(D)** Measurements of cell envelope thickness from cell membrane to cell wall edge were performed in FIJI as described in *Materials and Methods* with one measurement per individual cell indicated below the dot plot. Data are plotted as mean ± SEM from three biological replicates, with bars indicating the mean thickness and error bars indicating SEM. White arrowheads point to the membrane; black arrowheads point to cell wall, ****p<0.0001; ns, not significant; one-way ANOVA with Tukey’s post-test.

Because increased incorporation of staphyloxanthin in the membrane has been shown to rigidify the membrane (66, 67), we wondered if this increased staphyloxanthin incorporation was restraining SaeS kinase activity. To test this, we utilized an available transposon mutant in *crtM* encoding the 4,4’-diapophytoene synthase that is critical for staphyloxanthin synthesis (*crtM::tn*) and measured *sae*P1-*gfp* promoter activity in the *crtM::tn* single mutant and the Δ*pyrE crtM::tn* double mutant. We found that disrupting *crtM* had no effect on Sae activity in either genetic background, indicating that the increased staphyloxanthan accumulation did not explain the Sae signaling defect (**Fig. S7B**). Rather, the increased staphyloxanthin production may be in response to the irregularities we observe in the cell membrane, and an attempt to reinforce an unstable membrane. Taken together, these data suggest that pyrimidine sufficiency is correlated with proper cell envelope biogenesis.

### Pyrimidine sufficiency promotes SaeS kinase activity by reducing free fatty acid levels in the membrane

Cytidine nucleosides play a central role in lipid synthesis in prokaryotes and eukaryotes (68). Phosphatidate cytidylyltransferase (CdsA) condenses phosphatidic acid (PA) with CTP to form activated diacylglycerol (CDP-diacylglycerol; CDP-DAG) with the elimination of pyrophosphate. Displacement of CMP and cleavage of the phosphate monoester yields phosphatidylglycerol (69, 70). Previous studies have reported that the buildup of CDP-DAG precursors, specifically unincorporated free fatty acids (FFAs) in *S. aureus* membranes, is shown to inhibit the Sae TCS (71, 72). We reasoned that low CTP levels during pyrimidine limitation would reduce CdsA activity and lead to increased incorporation of FFAs into the membrane. To test this, we grew WT and Δ*pyrE* mutant cells to stationary phase (16 h) in the presence of fatty acid-free bovine serum albumin (BSA) and measured *sae*P1 promoter activity. BSA has been shown to bind and remove FFAs from the cell membrane, providing an extracellular sink for these lipids and reviving Sae TCS activity (43, 71–73). As expected, BSA treatment resulted in a modest increase in Sae activity in WT cells. Surprisingly, BSA treatment restored Sae activity to WT levels in the *ΔpyrE* mutant (**Fig. 5A**). We note that BSA addition did not restore pyrimidine auxotrophy of the mutant when grown in CDM (data not shown). A two-fold increase in BSA concentration had no further effect on Sae activity (**Fig. S8A**). LC-MS/MS analysis of supernatant samples revealed differences in the abundances of specific FFAs recovered using BSA treatment (**Fig. 5B**). Interestingly, we noticed that there was a dramatic increase in the abundance of 13:0 FFAs and a moderate increase in the abundance of 20:0, 22:0, and 24:0 FFAs recovered from the media from WT and *ΔpyrE* mutant cultures treated with BSA, where Sae activity is restored, relative to the untreated controls (**Fig. 5AB**). Of the FFAs identified, the 13:0 species seemed to be the highest FA in abundance in the media from both the WT and Δ*pyrE* mutant cells treated with BSA. However, it is unclear if 13:0 is limiting Sae activity. Even though LC-MS/MS allows us to identify FFA species, we cannot distinguish between different isoforms of the fatty acids detected. Given that anteiso branched-chain fatty acids derived from isoleucine (*a*15:0) promotes Sae activity (43, 44), we wondered if adding this fatty acid would rescue Sae activity in the Δ*pyrE* mutant as it is dramatically reduced in the supernatants of *ΔpyrE* mutant cultures during limitation. As expected, exogenous *a*15:0 restored Sae activity in a strain depleted for this membrane fatty acid (*lpdA::tn mbcS1*) (43, 44, 74). To our surprise, the addition of exogenous *a*15:0 led to a further reduction of Sae activity in the Δ*pyrE* mutant cells (**Fig. S8B**). There was no additional effect of *a*15:0 addition on growth. These data support the idea that the SaeRS defect seen in Δ*pyrE* mutant cells stems from FFA accumulation in the membrane due to a pyrimidine limitation-induced bottleneck during phospholipid biosynthesis.

**Figure 5:**
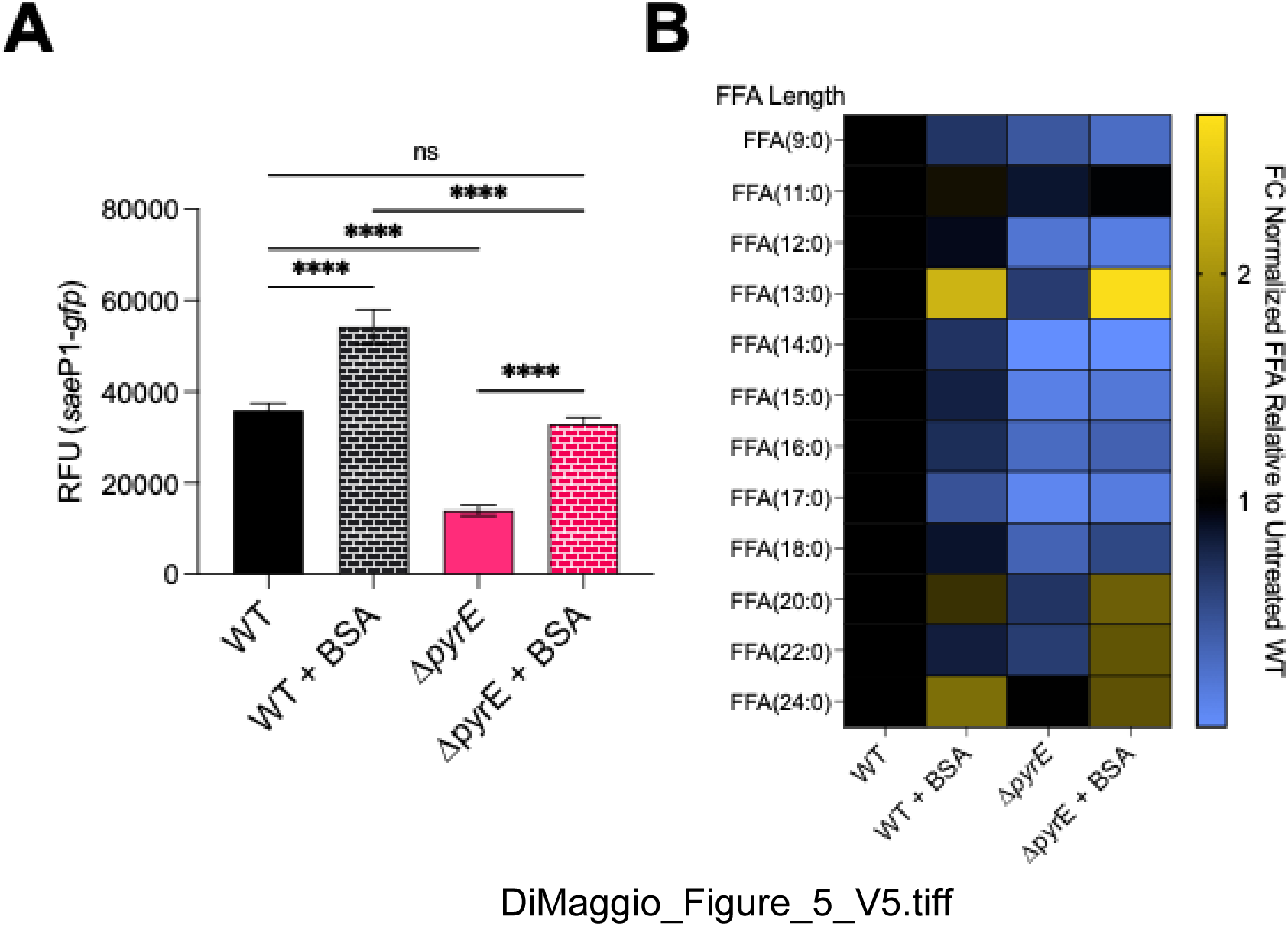
Treatment with bovine serum albumin restores Sae activity and promotes release of free fatty acids during pyrimidine limitation. (**A**) The indicated strains carrying the *sae*P1-*gfp* reporter were grown in TSB ± BSA (10 mg/mL^-1^) to stationary phase, at which time *sae*P1 promoter activity was measured. Data are plotted as mean ± SEM. (B) Culture supernatants from the indicated strains were collected, FFAs were extracted, and extracts were subjected to LC-MS/MS analysis. Data were normalized using an internal standard (d4-Palmitic acid) and mean fold change (FC) in abundances of free fatty acids (FFA) relative to WT are depicted using a heat map. All data are from at least three biological replicates ****p<0.0001; ns, not significant; One-way ANOVA with Tukey’s post-test.

## Discussion

This study aligns with the idea that nucleotide metabolism is an important factor for regulating virulence in *S. aureus* among other pathogens. Previous work from our lab and others have shown that mutations in *de novo* purine biosynthesis are correlated with altered expression of the virulence regulatory network (75–77). Specifically, mutations in the GMP synthase (GuaA) result in increased activity of the AgrCA TCS and enhanced protease activity *in vitro*. Further, it has been shown that the purine biosynthesis regulatory protein, PurR, can moonlight as a virulence regulator and represses expression of fibronectin binding proteins and extracellular toxins *in vivo* and *in vitro* (75, 78). On the contrary, mutations in *de novo* pyrimidine biosynthesis are correlated with a reduction in expression of virulence factors *in vitro* and *in vivo*. One study specifically looked at the impact of a mutation in the thymidylate synthase, ThyA, which was which downstream of the *de novo* pathway and is responsible for the conversion of deoxy uridine monophosphate (dUMP) into thymidylate (dTMP) (11).

They report that defective thymidylate synthesis is associated with a dramatic reduction *S. aureus* virulence *in vitro* and *in vivo* (11). Interestingly, regardless of the effects of purine and pyrimidine metabolism on enhancing or reducing virulence *in vitro*, *S. aureus* must be able to synthesize nucleotides *de novo* to support and maintain proper envelope homeostasis and the various cellular processes for proliferation within the otherwise nucleotide desert that is the host environment. This dual role of nucleotide metabolism in supporting both staphylococcal fitness and virulence makes it particularly attractive for the development of novel therapeutics. Thus, using an inhibitor that targets the *de novo* purine or pyrimidine biosynthetic would significantly hinder bacterial fitness and put the bacterium in a situation where the virulence regulatory systems could not function properly, ultimately leading to host clearance. Although purine and pyrimidine metabolism have been considered determinants for virulence potential, there is little mechanistic information about how these critical pathways contribute to virulence regulation. Here, we propose a model where pyrimidines act indirectly on the activity of the SaeRS TCS in *S. aureus* by maintaining membrane homeostasis (**Fig. 6**).

**Figure 6:**
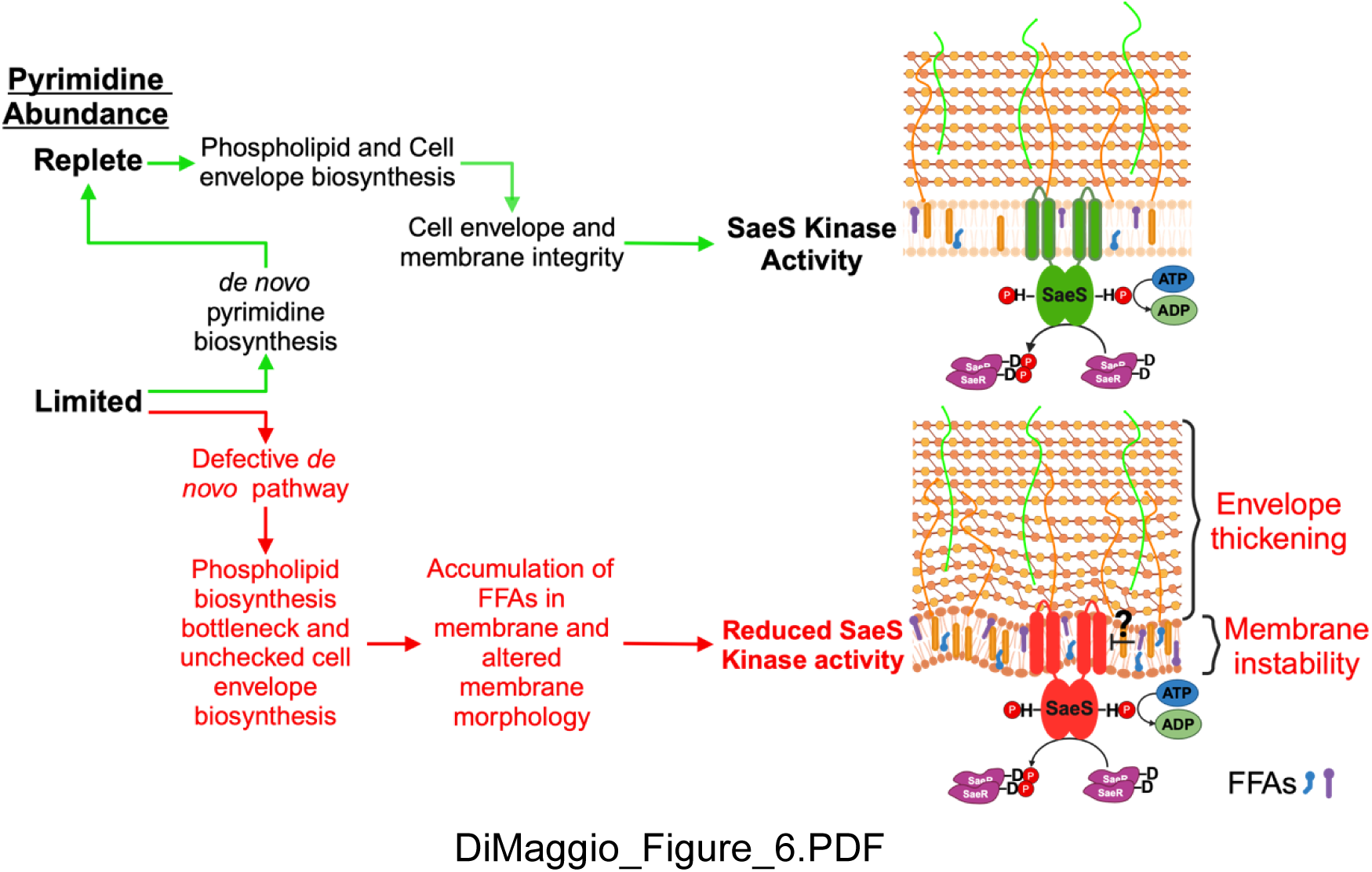
Working model for proposed mechanism by which pyrimidines promote SaeS kinase activity. During cellular growth, pyrimidine sufficiency dictates whether *S. aureus* needs to turn on the *de novo* pyrimidine biosynthetic pathway. Pyrimidines promote SaeS kinase activity, in part, by promoting and maintaining cell envelope and membrane integrity. Under pyrimidine-limited conditions where *S. aureus* cannot synthesize pyrimidines *de novo*, cell envelope biosynthesis goes unchecked, resulting in increased LTA production, thickening of the cell wall, and formation of a bottleneck in phospholipid biosynthesis. This results in an unstable membrane due to accumulation of phospholipid precursors, including free fatty acids, that act to reduce SaeS kinase activity.

Moreover, this study provides a unique perspective on the importance of membrane homeostasis and offers insight into how these IM-HKs, such as SaeS, are regulated.

To determine the mechanistic basis for the role of pyrimidine sufficiency in regulating the activity of the SaeRS TCS, the first task was to distinguish between metabolite accumulation and pyrimidine limitation as the root cause. Based on what is known about the role of orotate in regulating virulence in other organisms (15, 79), we initially hypothesized orotate or other pyrimidine metabolites were antagonizing Sae activity.

Although we observed orotate crystal formation in the *ΔpyrE* mutant colonies on solid medium, we determined that the defect was not due to orotate accumulation as the *ΔpyrD* and *ΔpyrC* mutants, which do not accumulate orotate and dihydroorotate, phenocopy the Δ*pyrE* mutant (**Fig. 2A**). Further, Sae activity is unaffected in pyrimidines replete conditions, consistent with the hypothesis that the defect is caused by pyrimidine limitation (**Fig. 2B**). We note that we cannot exclude the possibility that providing exogenous pyrimidines represses the *pyr* genes, preventing the accumulation of an inhibitory metabolite up stream of PyrC in the pathway. Moreover, it has been shown that purine and pyrimidine metabolism directly influence the activity of their respective pathways primarily through feedback inhibition of the regulatory proteins, PurR and PyrR. Although these two proteins have distinct regulatory mechanisms, they both function by sensing final or intermediate products of the pathways to control the expression of genes involved in nucleotide biosynthesis, thereby maintaining nucleotide balance within the cell (80). Given that feedback inhibition occurs, we were not surprised to see that most genes involved in purine biosynthesis were downregulated, while the genes encoding putative proteins involved in the *de novo* pyrimidine biosynthetic pathway (*pyrB*, *pyrC*, and *pyrD*) were upregulated in the Δ*pyrE* mutant during pyrimidine limitation (**Table 2; Table S2; Fig. 1A**). This underscores the influence that each respective biosynthetic pathway has on one another.

Our findings suggest pyrimidines affect SaeS through its N-terminal membrane-spanning domain, which is consistent with previous work showing IM-HKs like SaeS are sensitive to membrane composition changes (43, 44, 71, 72, 81). This is evidenced by the absence of pyrimidine-dependent Sae activity defects when complementing a *Δsae ΔpyrE* mutant with a truncated SaeS lacking its membrane-spanning domain (**Fig. 3 and S4**). In addition to identifying the connection of the Sae phenotype to the membrane, this experiment ruled out the possibility that reduced ATP pools were to blame for the low Sae activity. We also note that the expression of SaeR in the *Δsae ΔpyrE* mutant cells complemented with the *sae*P3-*saeRSc* construct was enhanced relative to the new WT control (*Δsae* + *RSc*). Several possibilities could explain this unexpected result. First, the difference in expression may stem in issues with ectopic expression of these constructs at a different locus. Second, this may be due to increased stability of SaeR in the *ΔpyrE* mutant background, possibly through decreased degradation by reduced activity of the Clp proteases. Third, the single mutant expressing the *sae*P3-*saeRSc* construct is the anomalous result, and the heightened activity seen in the Δ*pyrE* mutant is the expected result based on previous work with showing heightened Sae activity in the otherwise WT background (40).

Regardless of expression and stability of the proteins, there is still a dramatic enhancement in the overall kinase activity of SaeSc when expressed in the Δ*pyrE* mutant background (**Fig. S4**). Further work is needed to dissect the root cause of this difference in expression profiles of the P3*saeRSc* construct.

We observed dramatic changes to the cell wall architecture when pyrimidines are limited. Our observations are consistent with other reports that show defects in central metabolism result in an altered cell envelope (11, 43, 44, 61, 74, 77). Since the MICs for envelope-targeting drugs remained largely unchanged beyond the effects on LTAs in ΔpyrE mutant cells under limitation, and the SaeS kinase defect is linked to its N-terminal membrane domain, our findings suggest that envelope alterations create an environment unfavorable for full SaeS kinase activity. Previous studies showed that unchecked LTA synthesis results in a virulence defect but did not measure the activities of the major virulence regulatory systems AgrCA and SaeRS (64, 65). Both the LTA and phospholipid biosynthetic pathways share diacylglycerol (DG) as a common intermediate. Our data suggest that the accumulation of LTA in the *ΔpyrE* mutant may be the result of diverting metabolic flux from phospholipid synthesis to LTA synthesis (**Fig. S5)**(82). Accumulation of LTA in the *ΔpyrE* mutant background may help the cells compensate for the defect in phospholipid biosynthesis as LTAs serve important roles in processes associated with the envelope including ion homeostasis, maintaining proton motive force, as well as regulating autolysin activity (83, 84). In this study we also discovered that increased LTA synthesis is associated with a defect in SaeS kinase activity. This is consistent with a prior study that reported the link between *mspA* and Sae-dependent virulence factors (65). However, during pyrimidine limitation, increased LTA synthesis does not seem to be responsible for the Sae defect. Our observation that LTAs affect Sae is nevertheless important to note. How might this work? It is conceivable that SaeS may sense changes in envelope charge. LTAs are decorated with various negatively charged sugars. Incorporating D-alanine allows the bacterium to modify the envelope charge as a way to protect itself from host-mediated killing, resulting in a more positively charged envelope (63, 82). Increased LTA incorporation might alter envelope charge if D-alanylation machinery (*dlt*) cannot accommodate the higher LTA flux, thus creating an unfavorable environment for maintaining SaeS activity (83, 84). If so, overexpressing either *mspA* or the *dlt* may rescue of Sae activity when LTA production and/or envelope charge is altered. However, more work is needed to dissect the connection between LTAs and SaeS.

## Materials and Methods

### Bacterial strains and growth conditions

*Staphylococcus aureus* strains used in this study are listed in **Table S4**. Unless otherwise noted, indicated *S. aureus* strains were cultured in tryptic soy broth (TSB) containing 0.25% (wt/vol) dextrose (BD Biosciences) or chemically defined medium (CDM, pH 6.5) lacking pyrimidine nucleotides as previously described. Briefly, CDM medium was formulated with alanine (672 μM), arginine (287 μM), aspartic acid (684 μM), cysteine (166 μM), glutamic acid (680 μM), glycine (670 μM), histidine (129 μM), isoleucine (228 μM), leucine (684 μM), lysine (342 μM), methionine (20 μM), phenylalanine (240 μM), proline (690 μM), serine (285 μM), threonine (260 μM), tryptophan (50 μM), tyrosine (275 μM), valine (684 μM), thiamine (56 μM), nicotinic acid (10 μM), biotin (0.04 μM), pantothenic acid (2.3 μM), MgCl2 (1,000 μM), CaCl2 (100 μM), monopotassium phosphate (40,000 μM), dipotassium phosphate (14,700 μM), sodium citrate dehydrate (1,400 μM), magnesium sulfate (400 μM), ammonium sulfate (7,600 μM), and glucose (27,753 μM) (85). *Escherichia coli* strains were grown in lysogeny broth (LB) without glucose (10 g/L tryptone, 5 g/L yeast extract, and 5 g/L sodium chloride)(86). When necessary, media were solidified with agar (1.5% [wt/vol]) and supplemented with antibiotics or chemicals at the indicated concentrations: ampicillin (Amp), 100 µg/mL, chloramphenicol (Cm), 5 to 10 µg/mL; erythromycin (Erm), 5 µg/ml; tetracycline (Tc), 1.5 µg/mL; kanamycin (Km), 100 µg/mL; Bovine Serum Albumin (BSA), 10 mg/mL. When indicated, media were supplemented with exogenous pyrimidine (Uracil) to a final concentration of 200 µM. Unless otherwise noted, all strains were grown at 37°C.

I. **TSB growth curve:** Un-supplemented overnight cultures were used to inoculate 125-mL Delong flasks containing 12.5 mL TSB ± uracil supplementation (200 µM; 10:1 flask: medium ratio) to an optical density at 600 nm (OD_600_) of 0.05 and incubated in a water bath shaking at 250 rpm. Cell growth was monitored over 1 h increments by measuring increase in OD_600_ using an Amersham Ultraspec 2100 Pro UV-visible spectrophotometer.
II. **CDM grown curve:** Overnight cultures grown in CDM ± uracil supplementation (200 µM) were used to inoculate a 96-well plate to an OD_600_ of 0.05. To prevent uracil carryover from overnight cultures, 1 mL of cells were pelleted (13,000 x g for 1 min) and supernatants were removed. Cell pellets were washed twice with 1 mL of un-supplemented CDM before resuspension in 1 mL of un-supplemented CDM. Samples were then inoculated into fresh CDM medium ± uracil supplementation and cell density was monitored over time using a computer-controlled BioTek Synergy H1 plate reader (BioTek/Aligent) running Gen5 software v3.14.

### Genetic techniques

Oligonucleotides used for this study were synthesized by Integrated DNA Technologies (IDT; Coralville, IA) and are listed in **Table S5**. Restriction enzymes, DNA ligases, Exonucleases, DNA polymerases, Genomic DNA isolation, and PCR cleanup kits were purchased from New England Biolabs (NEB). Plasmids were extracted from *E. coli* using the Zyppy Plasmid Miniprep kit (Zymo). Unless indicated, plasmid constructions were performed using Gibson Assembly as described previously (87, 88). The plasmids used in this study are listed in **Table S6**. *E. coli* NEB 5α (NEB) was used as host for plasmid constructions and all plasmids were confirmed by PCR and plasmid sequencing (PlasmidSaurus) before being introduced into *S. aureus* strain RN4220 by electroporation (89). Plasmid and marked mutations were moved between *S. aureus* strains via Φ85-mediated mediated transduction(90).

### Plasmid Constructions

(i) **pDD02.** 500 bp upstream and downstream of *pyrE* was amplified from wild-type gDNA using the following primer pairs: oDD062, oDD063 (500 upstream), and oDD64, oDD65 (500 downstream) with Q5 DNA polymerase (NEB). pKOR1 (10,374 bb) was linearized and amplified using primers oDD60 and oDD61 (91). Each primer was designed to add 25 bp overlapping for later Gibson assembly. The linearized vector was treated with DpnI (NEB) for 1 h at 37°C and cleaned up using a PCR purification kit (NEB). The purified PCR products were then subjected to Gibson Assembly and then transformed into *E. coli* DH5a. **pDD03 and pDD10** were constructed in a similar manner for *pyrD* and *pyrC*.
(ii) **pDD05.** *pyrD* including its native promoter region were amplified from wild-type gDNA with the forward primer containing a SacI cutting site and the reverse primer containing a BamHI cutting site. Both the vector backbone, pLL39, and the *pyrD* insert were cut with BamHI and SacI (NEB) and treated with Quick CIP (NEB) to dephosphorylate the ends to prevent self-ligation. After CIP treatment, the products were cleaned up using a PCR purification kit (NEB). The purified linearized vector and insert were ligated using T4 DNA ligase (NEB) and the resulting plasmid was then transformed into *E. coli* DH5a.

### Construction of in-frame deletions

Plasmids pDD02 (Δ*pyrE*), pDD03 (Δ*pyrD*), and pDD10 (Δ*pyrC*) were moved to *S. aureus* strain LAC at the permissive temperature (30°C). Allelic exchange was performed as described using *secY* antisense counterselection (91). In-frame deletion alleles were verified by PCR and Sanger sequencing.

### GFP reporter assays

Cells carrying reporter fusions were streaked out from frozen stocks on TSA plates containing appropriate antibiotics. Single colonies were used to inoculate TSB with or without supplements: 200 µM uracil, 200 µM cytidine, 10 mg/mL fatty-acid-free bovine serum albumin (BSA). For growth in TSB ± uracil or cytidine, cells were grown in 3 mL medium in disposable 16×125 mm borosilicate glass test tubes. For BSA treatment, cells were grown with the 10:1 flask:medium ratio in DeLong flasks to stationary phase (16 h). Resulting cultures were then diluted 1:5 in 1X phosphate-buffered saline (PBS) to a total volume of 100 µL per well in a 96-well plate. Both OD_600_ and GFP fluorescence (485 nm excitation, 535 nm emission) were read in a computer-controlled BioTek Synergy H1 plate reader (BioTek/Aligent) running Gen5 software v3.14. Relative Fluorescent Units (RFU; GFP fluorescence/OD_600_) were calculated by subtracting the background fluorescence of TSB and dividing by OD_600_ to correct for cell density.

### Analysis of SaeS kinase activity

Separation of SaeR and SaeR∼P was performed as described previously (44) using 12% polyacrylamide gels containing 100 µM manganese and 50 µM of the acrylamide-pendant Phos-tag ligand (Wako Laboratory Chemicals). Cells were grown in TSB to mid exponential phase, early stationary phase and late stationary phase (2 h, 8 h, and 18 h, respectively) at 37°C at 10:1 flask to medium ratio with shaking. Cell pellets (OD_600_ of 10) were collected at 13,000 x g and stored at –80°C prior to analysis. Whole cell extracts were obtained by resuspending cell pellets in 300 µL cell extract buffer (20 mM Tris [pH 7.0], 1X Protease Inhibitor Cocktail Set I (Sigma-Aldrich)) and transferred to sterile screw cap tubes containing approximately 100 µL of 0.1 mm silica beads. The cells were homogenized at room temperature using a Precellys 24 bead beater (Bertin technologies) for 3 cycles of 6500 rpm for 30 s each, followed by 3 min pauses on ice.

The tubes were then centrifuged at 8500 x g for 15 s settle the beads, and the supernatant was transferred into new tubes. Whole-cell extracts were normalized by protein concentration (absorbance at 280 nm [A280]) to 100 µg and electrophoresed on Phos-tag gels with standard running buffer (0.1% [wt/vol] SDS, 25 mM Tris-HCl pH 8.3, 192 mM glycine) at 4°C under constant voltage (150 V) for 2 h. The gels were washed for 15 min with transfer buffer (25 mM Tris [pH 8.3], 192 mM glycine, 20% methanol) with 1 mM EDTA followed by a second wash without EDTA to remove manganese ions. Proteins were then transferred to 0.45 µM PVDF membranes (Cytiva). Membranes were blocked in blocking buffer (5% [wt/vol] skim milk in Tris-Buffered Saline with Tween 20 (TBST) (20 mM Tris-HCl pH 7.6, 150 mM NaCl with 0.1% [wt/vol] Tween 20) for 1 h.

Membranes were then subjected to three brief washes in TBST and incubated with polyclonal rabbit antibodies to SaeR (1.5:1,000) for 1 h. Membranes were then washed three times with and incubated with StarBright Blue 700 goat anti-rabbit IgG (1:3500; Bio-Rad) for 1 h. Membranes were subjected to three brief washes in TBST and signals were visualized using an Amersham ImageQuant800. The densities of the SaeR∼P relative to total SaeR signal were determined by quantification with Multi Gauge software (FujiFilm). The data are representative of three different independent experiments, and a representative image is shown.

### Analysis of SaeR protein levels

The whole cell extracts, and protein concentrations described above were used to visualize total SaeR levels. Briefly, 100µg of whole cell extracts were subjected to 12% SDS-PAGE, and proteins were transferred to 0.45 µM PVDF membranes (Cytiva). After transfer, membranes were blocked in blocking buffer for 1 h. Membranes were then subjected to three brief washes in TBST and incubated with polyclonal rabbit antibodies to SaeR (1.5:1,000) for 1 h. Membranes were then washed three times with and incubated with StarBright Blue 700 goat anti-rabbit IgG (1:3500; Bio-Rad) for 1 h.

Membranes were subjected to three brief washes in TBST and signals were visualized using an Amersham ImageQuant800. The densities of the SaeR signal were determined by quantification with Multi Gauge software (FujiFilm). The data are representative of three different independent experiments, and a representative image is shown.

### RNA Sequencing

Overnight cultures of cells grown in TSB were used to inoculate 125 ml DeLong flasks containing TSB with a 10:1 flask:media ratio. Flasks were incubated in a shaking water bath set to 37°C at 250 RPM for 8 h. 2 mL cells were collected at 13,000 x g, and resuspended in an equal volume of quenching solution (Ethanol:Acetone [1:1]), pelleted, and stored at –80°C until use. Cell pellets were resuspended in TRIzol and transferred to sterile screw cap tubes containing approximately 100 µL of 0.1mm silica beads. The cells were then homogenized at room temperature using a Precellys 24 bead beater (Bertin technologies) for 3 cycles of 6800 rpm for 30 s each, followed by 5 mins pauses on ice. The tubes were then centrifuged at 4 °C and 13,000 x g for 1 minute to settle the beads. Clarified extracts were then mixed in an equivalent volume of 100% [v/v] ethanol. RNAs were purified using a Direct-Zol RNA Miniprep kit (Zymo Research) according to manufacturer’s instructions and RNA yields were determined by measuring (A260/280). RNA quality was determined using a 5200 Fragment Analyzer System (Aligent) and RNA-Seq libraries were generated using an Illumina Stranded Total RNA kit after rRNA depletion (Ribo Zero Gold; Illumina). The resulting RNA-Seq libraries were then sequenced on a HiSeq 2500 Sequencing System (Illumina).

### Differential Expression Analysis

The sequencing files were processed with Trimmomatic (92). Alignment to the *S. aureus* USA300_FPR3757 genome (NC007793; NCBI) and pairwise comparisons were made using Rockhopper (Wellesley College) (93, 94). Raw RNA-seq datasets were uplo

### KEGG Analysis

Outputs for RNA-Seq were then processed using the KEGGREST package in R to determine which biological pathways were enriched or disturbed based on the KEGG input genome for the USA300_FPR3757 isolate (saa; KEGG) (53, 54).

### Antibiotic Minimum Inhibitory Concentration (MIC) measurements

MIC measurements were performed according to the Clinical and Laboratory Standards Institute (CLSI) methods for broth microdilution susceptibility testing of staphylococci (59). Strains were grown on cation-adjusted Mueller Hinton broth (caMHB) plates containing 2% NaCl at 37°C for 24 h and 5-10 colonies were resuspended in 0.85% saline before adjusting to McFarland standard 0.5 (A_600_=0.1). The cell suspensions were then diluted 1:20 in sterile 1X PBS and 300 µl was used to inoculate 3 ml media containing serially diluted antibiotics or compounds (Tunicamycin, Fosfomycin, Oxacillin, Congo Red). The tubes were incubated at 35°C for 24 h and MIC values were recorded as the lowest compound concentration where no growth was observed.

### Lipoteichoic acid extraction and blotting

I. **Extraction:** Lipoteichoic acid extraction was performed as previously described (61, 64, 95). Single colonies grown on TSA plates were used to inoculate 3 mL cultures in disposable 16×125 mm borosilicate glass tubes (Fisher Scientific). Cells were grown to stationary phase overnight (16 h) at 37°C and incubated with rotation. After incubation, cells were normalized to an OD_600_ of 3 and collected at 7800 rpm. The supernatant was removed, and pellets were resuspended in 1 mL sterile 1X PBS and transferred to screw cap tubes containing approximately 100 µL of 100µm glass beads. The cells were homogenized at room temperature using a FastPrep-24 (MP Biomedicals) for 3 cycles of 6 m/s for 40 s each, followed by a 4-minute rest. Tubes were then centrifuged at 200 x g for 1 min to settle the beads and 700 µLof the supernatant was transferred into a fresh 1.5 mL Eppendorf tube. The tubes containing the supernatant were then centrifuged at 14,000 rpm for 15 min to collect the cell debris. After centrifugation, the supernatant was removed, and the remaining cell debris pellet was resuspended in 80 µL 2X Laemmli sample buffer (BioRad) and incubated at 95°C for 20 min. After heating, samples were centrifuged at 14,000 rpm for 5 min, and the supernatant was stored at –20°C until ready for western analysis.
II. **Blotting:** 20 µL of the LTA extracts were run on 15% polyacrylamide gels at room temperature under constant voltage (100V) until the BPB reached the bottom of the gel and transferred to PVDF. After transfer, membranes were blocked in blocking buffer for 1 h. Membranes were then subjected to three brief washes in TBST and incubated with monoclonal mouse antibodies specific for LTAs (mAB55; HycultBiotech; 1:5000). Membranes were then washed three times with and incubated with StarBright Blue 700 goat anti-mouse IgG (1:3500; Bio-Rad) for 1 h. Membranes were subjected to three brief washes in TBST and signals were visualized using an Amersham ImageQuant800. The densities of the LTA signal were determined by quantification with Multi Gauge software (FujiFilm). The data are representative of three different independent experiments, and a representative image is shown.

### Staphyloxanthin analysis

The extraction of pigment from *S. aureus* was performed as previously described (96). Single colonies grown on TSA plates were used to inoculate 3 mL cultures in disposable 16×125 mm borosilicate glass tubes (Fisher Scientific). Cells were grown to stationary phase (24 h) at 37°C and incubated with rotation. After incubation, 1 mL of culture was transferred to a 1.5 mL microcentrifuge tube and pelleted at 13,000 x *g* for 1 min. The supernatant was removed, pellets were washed with sterile distilled water, followed by re-centrifugation and removal of supernatant. Cell pellets were then resuspended in 200 µl of methanol followed by incubation at 55°C for 3 min in an Eppendorf ThermoMixer.

After incubation, the cell-methanol mixture was centrifuged at 13,000 x *g* to remove cell debris, and the supernatant containing the pigment was transferred to a new microfuge tube. To ensure total removal of pigment, the extraction was repeated, and the extracts were pooled together. 100 µl of extracts were then transferred to quartz cuvettes containing 900 µl methanol and mixed thoroughly by pipetting before measuring at 465 nm in an Amersham Ultraspec 2100 Pro UV-visible spectrophotometer. Absorbance was normalized by CFU enumeration by plating serially diluted cells to TSA plates.

### Transmission Electron Microscopy

Single colonies grown on TSA plates were used to inoculate 4 mL of TSB ± uracil supplementation (200 µM). Cells were grown to stationary phase (16 h) at 37°C and incubated with rotation. After incubation, cells were pelleted in 2 mL microcentrifuge tubes until the pellet was ∼2 mm tall from the bottom of the tube. Cell pellets were then washed three times in sterile 1X PBS (pH 7.4). Samples were then fixed overnight at 4°C in sterile 1X PBS (pH 7.4) containing 2.5% [vol/vol] EM grade glutaraldehyde before being processed at the Laboratory of Biological Ultrastructure at the University of Maryland, College Park, MD. Subsequently, cells were subjected to a second fixing step in 1% [v/v] osmium tetroxide for 1 h. Samples were then washed 3 times in double distilled water and postfixed in 2% [v/v] uranyl acetate solution for 60 min. Samples were then dehydrated using five incremental ethanol washes (35%, 50%, 70%, 95%, 100% [v/v] for 10 min each wash); the final wash using ethanol (100% [v/v]) was repeated 3 times. Following dehydration, cells were infiltrated using propylene oxide (100% [v/v]) two times for 15 min each followed by overnight incubation with agitation in a 1:1 [v:v] mixture of propylene oxide (100% [v/v]) and resin (100% [v/v]). The following morning, samples were subjected to incremental infiltration steps (1:2, 1:3, 100% resin; propylene oxide (100% [v/v]): resin (100% [v/v]) for 60 min each step); the final infiltration step with 100% resin was repeated before embedding the samples in fresh resin. Samples were then solidified by incubating the resin at 70 °C for a total of 12 days before thin-sectioning and staining. Stained sections were loaded onto mesh grids for imaging. Images were obtained on a Hitachi HT7700 Transmission Electron Microscope at the Laboratory of Biological Ultrastructure at the University of Maryland, College Park, MD. Cell diameter and envelope thickness were measured in FIJI (97).

Measurements were determined by first setting pixel values to the length of the scale bar, followed by use of the line tool to measure cell diameter or envelope thickness (from cell membrane to the outside of the cell wall), respectively.

### Medium free fatty acid (FFA) analysis using QTRAP 5500

This method is designed to measure 21 classes of lipid molecules which includes free fatty acids (FFA) using QTRAP® 5500 LC-MS/MS System (Sciex). Single colonies were used to inoculate 125 mL DeLong flasks containing TSB ± BSA (10mg/mL) with a 10:1 flask:medium ratio. Flasks were incubated in a shaking water bath set to 37°C at 250 rpm for 16 h. Resulting cultures were checked for promoter activity then pelleted at 3500 x g for 20 min in a sterile 50 mL falcon tube. After pelleting the cells, 9 mL of the supernatant was transferred to a fresh 50mL falcon tube. For FFA analysis, 2 ml supernatant from resulting promoter activity cultures was freeze dried and used for sample preparation. 125 µL of pre-chilled isopropanol containing internal standard (FFA-d4-Palmitic acid – 1ug per mL; Cambridge Isotope Laboratories) was added and samples were vortexed. The sample tube was plunged into dry ice for 30 s followed by a 90 s incubation in a 37 °C water bath. The freeze thaw cycle was repeated for three times followed by sonication for 30 s. After sonication, samples were vortexed for 1 min and were incubated at –20°C for overnight for complete protein precipitation. The samples were then centrifuged at 13,000 rpm for 20 min at 4°C. After centrifugation, the supernatant of each sample was transferred to a MS vial for LC-MS analysis. 5 µL of each sample was injected onto a Xbridge amide 3.5 µm, 4.6 × 100 mm (waters) using SIL-30 AC auto sampler (Shimazdu) connected with a high flow LC-30AD solvent delivery unit (Shimazdu) and CBM-20A communication bus module (Shimazdu) online with QTRAP 5500 (Sciex, MA, USA) operating in positive and negative ion mode. A binary solvent comprising of acetonitrile/water 95/5 with 10 mM ammonium acetate as solvent A and acetonitrile/water 50/50 with 10 mM ammonium acetate as solvent B was used for the resolution. Lipids were resolved at 0.7 mL/min flow rate, initial gradient conditions started with 100% of solvent A at 0.2 ml per min flow rate, shifting towards 99.9% of solvent A over a time period of 3 min, 94% of solvent A over a time period of 2 min at 5 min and 75% of solvent A over a period of 2 min. Finally, equilibrating to initial conditions (100% of solvent A) over a time period of 3 minute using auto sampler temperature 15°C and oven temperature 35°C. Source and gas setting were used as follows: curtain gas = 30, CAD gas = medium, ion spray voltage = 5.5 kV in positive mode and −4.5 kV in negative mode, temperature = 550 °C, nebulizing gas = 50 and heater gas = 60. The data were then normalized to the respective internal standard area (FFA-d4-Palmitic acid; Cambridge Isotope Laboratories) and processed using MultiQuant 3.0.3 (Sciex). The quality and reproducibility of LC-MS data was ensured using several measures. The column was conditioned using the pooled QC samples initially and were also injected periodically to monitor shifts in signal intensities and retention time as measures of reproducibility and data quality of the LC-MS data.

### Statistical analysis

Data shown are the results of at least three independent experiments. Statistical significance was determined using Prism ver. 10 (GraphPad) with the indicated tests.

### Accession number

RNA-sequencing date have been deposited in the NCBI database under the BioProject accession number PRJNA1232367 and in the Sequence Read Archive (SRA) under accession number(s) (SRR32583580 through SRR32583585).

## Acknowledgements

We thank Timothy Maguel at the Laboratory of Biological Ultrastructure for technical assistance with TEM, Dr. Albert Tai and Irina Grinvald at Tufts University Core Facility (TUCF) Genomics Core for the support of the included NGS analysis, Dr. Taeok Bae (Indiana University School of Medicine-Northwest) for the gift of SaeR antibodies and pMA-15, Dr. Ruth Massey for the gift of the *mspA::tn* mutant, and Dr. Shivani Bansal in the Georgetown University Metabolomics and Proteomics Shared Resources for help with LC-MS/MS identification of FFA. We also thank Dr. Alexander Horswill (University of Colorado Anschutz School of Medicine), Brinsmade lab members, and other members of the Microbial Interest Group at Georgetown University for helpful comments and suggestions on the science reported here. This work was supported by NIH grants R01, R56 AI137403 to SRB. TUCF Genomics was supported by NIH grant 1S10OD032203-01 and Georgetown Metabolomics was partially supported by NIH/NCI/CCSG grant P30 CA051008-29S2. The funders had no role in study design, data collection and analysis, decision to publish, or preparation of the manuscript.

## Author Contributions

DAD and SRB conceived the study; WY originally identified *pyrE::tn* mutant; DAD and SRB conceptualized the research goals and aims; DAD performed the investigations and analyzed the data, DAD and SRB prepared the original manuscript draft; DAD, WY, and SRB prepared the final manuscript.

